# NRT1.1s and NRT2.1 affect rhizosphere and apoplastic pH during plant nitrate uptake

**DOI:** 10.1101/2025.10.30.685715

**Authors:** Qiying Xiao, Junjie Wang, Yi Chen, Fuyu Li, Wuge Jiwu, Anthony J. Miller, Jeremy D. Murray

**Affiliations:** Department of Botany, CAS-JIC Centre of Excellence for Plant and Microbial Science (CEPAMS), Centre for Excellence in Molecular Plant Sciences (CEMPS), Shanghai Institute of Plant Physiology and Ecology (SIPPE), Chinese Academy of Sciences, 300 Feng Lin Road, Shanghai 200032, China; Department Biochemistry & Metabolism, John Innes Centre, Norwich Research Park, Norwich, NR4 7UH, UK; College of Life Sciences, Shanghai Normal University, Shanghai, China

**Author notes:** **Corresponding author:** Jeremy D. Murray **Email:**.

**Keywords:** nitrate transport, apoplast, cell wall, pH

## Abstract

Nitrate is the primary source of nitrogen for plants in most environments. Nitrate uptake from the rhizosphere into root cells is mediated by Nitrate Transporter 2 (NRT2) and NRT1.1 proteins via proton symport, with the apoplast serving as an intermediate compartment. Using a simple root immersion system, we investigated the effects of NRT transporters on external media pH (pH_med_) and apoplastic pH (pH_apo_) in *M. truncatula* nitrate transporter mutants. Monitoring of pH_med_ showed that loss of NRT2.1 largely prevented nitrate induced alkalization of the media, while loss of NRT1.1A/B reduced media alkalization only during the initial stages of nitrate uptake, suggesting they have fundamentally different physiological roles. Use of the pH sensitive dye HPTS showed that the pH_apo_ of root cells initially decreased during nitrate uptake, and this required NRT1.1A/B. Unexpectedly, the observed changes in pH_med_ during nitrate uptake in WT and the mutants were not reflected in pH_apo_. To explain these discrepancies, we propose a model in which the depletion of protons in the media occurring during nitrate symport is offset by the buffering of protons in the cell wall and the activity of the H^+^-ATPase, allowing a lower pH_apo_ to be maintained. Such a mechanism would enable efficient nitrate uptake across a range of external pH values.

## Introduction

Nitrate uptake in plants depends on proton symport, which requires a supply of protons from the apoplast. This requirement is reflected in the apoplastic pH (pH_apo_), which typically ranges from 4.5-5.5(1). Nitrate symport results in alkalization of the rhizosphere which occurs through the consumption of protons by nitrate proton symporters with a stoichiometry of at least 2 H^+^ per nitrate absorbed (2). In *Arabidopsis thaliana*, this occurs through the activities of the high affinity nitrate uptake transporter AtNRT2.1 and the low or dual affinity transporter AtNRT1.1, also called Nitrate Transporter 1/Peptide Transporter 6.3 (3,4). AtNRT1.1 also plays an important role in nitrate sensing (5). Loss of AtNRT1.1 greatly diminishes nitrate-induced rhizosphere alkalization (6), but NRT2.1’s role in this phenomenon has not been studied. Despite it playing a critical role in nitrate uptake, the dynamics of pH_apo_ in response to nitrate remains uninvestigated. Here we addressed these questions in *Medicago truncatula* (Gaertn) using NRT2.1 and NRT1.1A/B mutants, the respective orthologs of Arabidopsis NRT2.1 and NRT1.1 (7).

## Results

To observe the effect of nitrate uptake on medium pH (pH_med_), *M. truncatula* seedlings were grown on agar plates containing nitrate-free Fåhraeus nutrient solution (FP) for one week. The seedlings were then transplanted into non-aerated liquid FP medium supplemented with different concentrations of KNO_3_ (initial pH_med_ of 6.0) with their roots fully submerged. In the absence of nitrate, the seedlings acidified the FP medium to at ∼4.0-4.2 by 24h (Fig. 1*A*). Treatment with high concentrations of nitrate (≥5mM) significantly lowered pH_med_ at 3 and 6h compared to the FP control by 0.2-0.4 pH units (Dunnett’s test, p-value <0.05; inset Fig. 1*A*), but this trend reversed at 24h with the nitrate containing samples becoming alkalized, with a pH_med_ ∼1 pH unit higher than the nitrate-free control, with even larger differences (∼1.5-2.2 pH units) occurring at later time points. In comparison, treatment with low nitrate (0.5 mM KNO_3_) had only a slight effect on pH_med_. These results are consistent with media alkalization by the seedlings being due to nitrate/H^+^ symport mediated depletion of protons.

**Figure 1.**
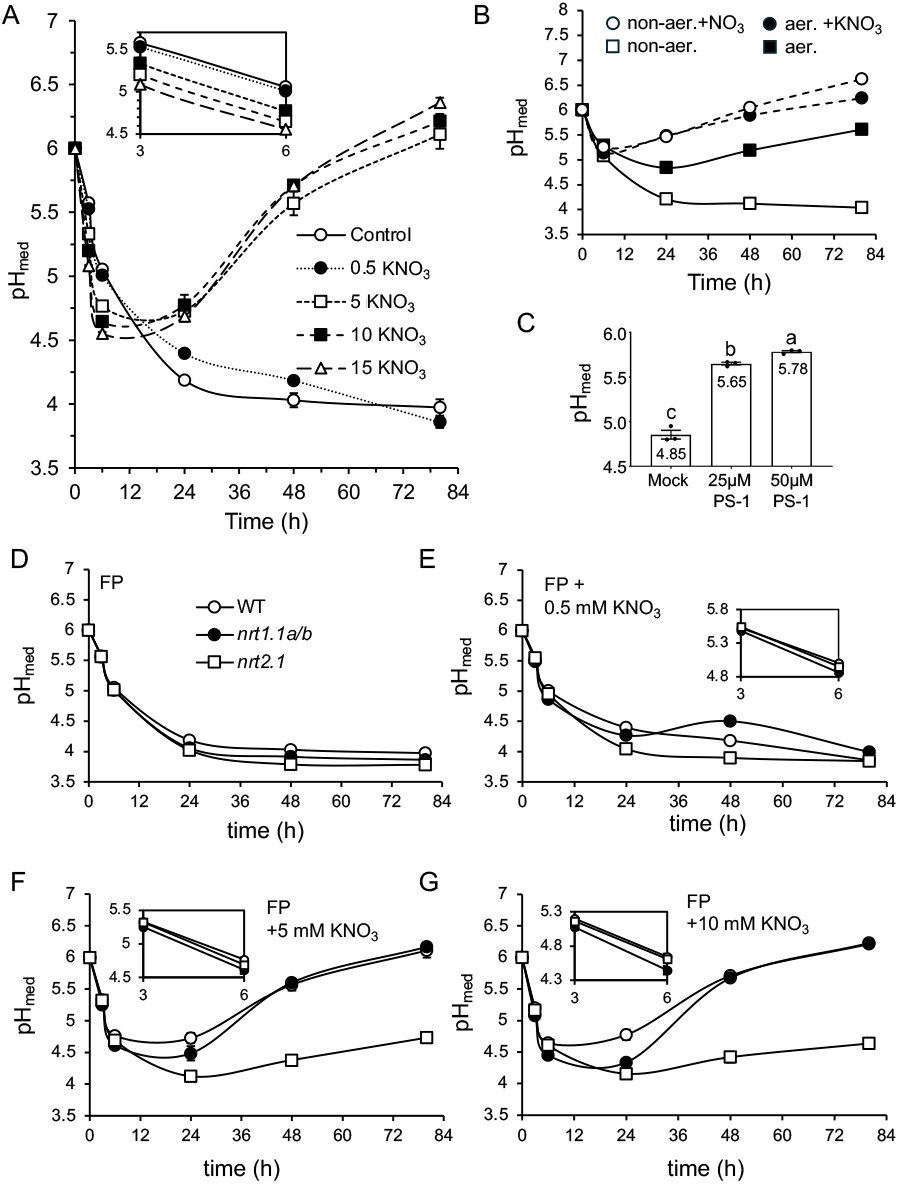
NRT2.1 and NRT1.1s affect nitrate-induced media alkalization. WT *M. truncatula* seedlings were transferred to either FP (control), or FP supplemented with varying concentrations of KNO_3_ and pH_med_ was monitored (*A*). *M. truncatula* seedlings were transferred to either FP or FP containing 5 mM KNO_3_ with and without aeration (aer.) and pH_med_ was monitored (*B*). Seedlings grown on FP were transferred to FP containing the indicated concentrations of PS-1 for 6h and then pH_med_ was determined (*C*). WT and mutant *M. truncatula* seedlings were transferred to either FP (*D*), or FP with varying concentrations of KNO_3_ (*E*-*G*) and pH_med_ was monitored. In (*A*), the inset shows a detailed view of the 3 and 6 h timepoints. For all experiments *N*=3, with 5 plants per biological replicate. Error bars indicate SEM.

To determine if the medium acidification in our system was caused by flood-induced hypoxia, we repeated the experiment with gentle aeration. This increased the O_2_ concentration of the media from 7.2 (±0.04 SEM) to 9.6 (±0.17) ppm and largely attenuated the acidification (Fig. 1*B*). We then investigated the potential involvement of the PM H^+^-ATPase using the specific inhibitor protonstatin-1 (PS-1). Treatment with PS-1 essentially blocked root proton extrusion, with 91 and 95% inhibition by 25 and 50mM PS-1, respectively (Fig. 1*C*). These results suggest that the observed media acidification is mediated by the PM H^+^-ATPase and is induced by short-term hypoxia resulting from the immersion.

The root immersion system provides a means of evaluating the H^+^ consumption by NRTs during nitrate uptake. First, we tested whether the acidification of the media was affected by the absence of NRTs. Five mutants were tested, *nrt2*.*1, nrt1*.*1a, nrt1*.*1b, nrt1*.*1c*, and the *nrt1*.*1a/b* double mutant. Every mutant acidified pH_med_ to ∼3.8-4.0 within 24h, similar to WT (Fig. 1*D*). We then tested the effect of these mutations on pH_med_ in the presence of various concentrations of KNO_3_. We found that the *nrt1*.*1a/b* double mutant and *nrt2*.*1* mutant affected pH_med_ in different ways (Fig. 1*E-G*). The *nrt1*.*1a/b* mutant showed a small but significant (∼0.1 pH unit; Dunnett’s test, p-value <0.05) decrease in pH_med_ at 3 and 6h after nitrate addition at all nitrate concentrations. This effect grew stronger at 24h (∼0.2-0.4 pH units, depending on the nitrate concentration) but was greatly diminished or absent at the later time points. This sharply contrasted with *nrt2*.*1*, which showed no phenotype at 3 and 6h, but had a significantly diminished ability to alkalize the media at 24 and 48 under all nitrate concentrations, which grew more pronounced at 80h (∼1.4-1.7 pH units) under 5 and 10 mM KNO_3_. Under 0.5mM the pH_med_ phenotype of *nrt2*. at 24h and 48h was weaker, and no difference was seen at 80h.

We then investigated whether changes occur in pH_apo_ occur during nitrate uptake. We used HPTS, a widely used fluorescent membrane-impermeant pH sensitive dye, to stain immersed roots and made observations on root tip cells using confocal microscopy (see *S1 Appendix*). After transfer to 5 mM nitrate the pH_apo_ of WT seedlings significantly decreased, indicating a pH drop, at 4h (Fig. 2*A*). The *nrt1*.*1a/b* mutant failed to respond, indicating it has a reduced ability to acidify the apoplast. Observations at the 24- and 48-hour marks indicate that pH_apo_ of WT recovered, while that of *nrt1*.*1a/b* was higher at 48h.

**Figure 2.**
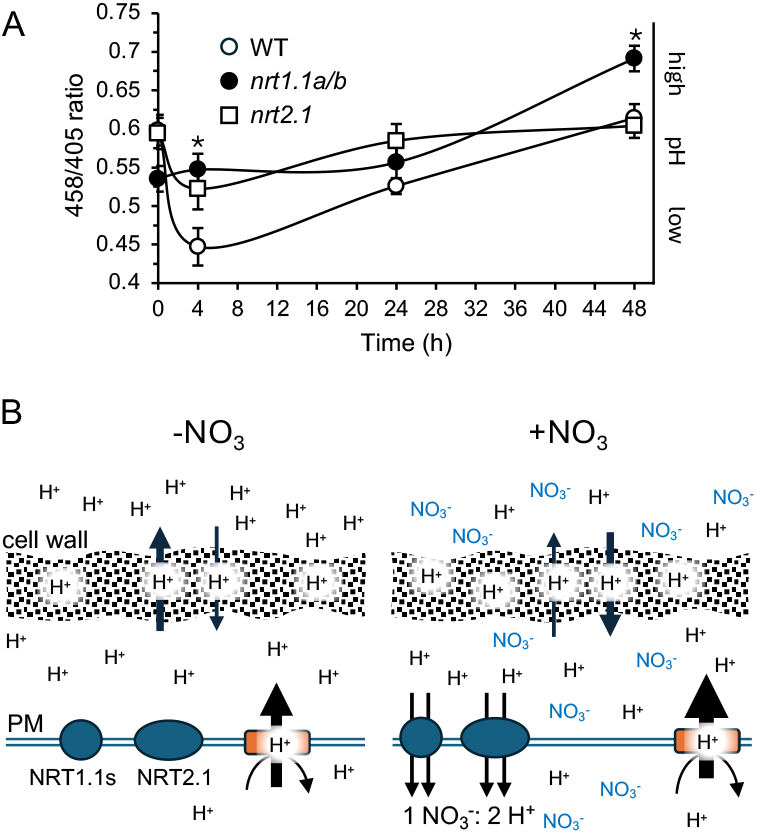
Nitrate induces a short-term decrease in pH_apo_ which requires NRT1.1A/B. Roots of *M. truncatula* seedlings were stained with HPTS and 458/405 ratios of the root tip cells were determined by confocal microscopy after immersion in FP (0 h) or at different times after addition of FP containing 5mM KNO_3_ (*A*). *P-value<0.05, Dunnett’s test, mutants vs WT at each time point. *N*=6-32. A model for nitrate uptake in plants (*B*). (left) Without aeration and in the absence of nitrate the rhizosphere and apoplast of immersed plants are acidified through the activity of H^+^-ATPases that serve to maintain cytosolic pH during hypoxia. (right) When nitrate is added to the media, the activation of NRT1.1 and NRT2.1 symporters triggers an overcompensation of the PM H^+^-ATPase through an unknown mechanism, which more than offsets H^+^ uptake via nitrate symport, leading to a small decrease in pH_apo_. The ongoing uptake of nitrate and protons via the apoplast by the symporters eventually leads to the alkalization of the rhizosphere.

## Discussion

The seedling immersion approach provides a convenient means to observe the effect of root nitrate uptake on external pH. Our results indicate that the initial drop in pH_med_ after seedling transfer was caused by hypoxia. A similar drop in pH_med_ was reported in *A. thaliana* after root immersion, which was shown to require H^+^-ATPases 1 and 2 (8). Furthermore, it was shown that hypoxia induces increased proton efflux in barley root tips, which was attributed to PM H^+^-ATPase activity (9), suggesting this is a general hypoxia-related phenomenon. Using this approach, we found that nitrate initially induced a small drop in pH_med_ and pH_apo_. It was shown in maize that following an initial depolarization after exposure to nitrate, the PM becomes hyperpolarized through overcompensation of the H^+^-ATPase (10). Such hyperpolarisation could explain the observed drop in pH_med_ and pH_apo_ at early time points. Consistent with this, reports in maize showed an increase in root PM H^+^-ATPase activity after nitrate supply (11,12).

The initial nitrate-induced drop in pH_apo_ was compromised in the *nrt1*.*1a*/*b* mutant. This requires an explanation, since decreased nitrate/H^+^ symport would be expected to lower, not raise, the pH of this compartment. We suggest that this may occur through reduced activation of the PM H^+^-ATPase due to lower nitrate uptake in the mutants. While the trigger for nitrate induced hyperpolarization is unknown, it is possible that the increased influx of protons into the cytoplasm by nitrate/H^+^ symport could trigger higher H^+^-ATPase activity to maintain a stable cytoplasmic pH (13). In this case, a decrease in nitrate/H^+^ symport activity would lessen the stimulation of the PM H^+^-ATPase by alleviating symport-mediated cytosolic acidification, resulting in a higher pH_apo_ in the mutants. Another, non-exclusive possibility, is the inhibitory effect of nitrate on vacuolar (V)-ATPases (14). Such downregulation of the V-ATPase would serve a complementary role, preventing competition with the PM-ATPase for cytosolic protons during nitrate uptake.

The strong alkalization observed in the medium at later time points did not occur in the apoplast. Similarly, the much lower pH_med_ of *nrt2*.*1* compared to WT at 48h was not reflected in its pH_apo_, which was similar to WT. It is generally assumed that pH_apo_ and pH_med_ are the same since these compartments are physically connected. These discrepancies can be explained by the role the cell wall plays in maintaining a low pH_apo_. The cell wall’s apoplast has two zones: the Water Free Space, which mirrors the ionic concentration of the external solution (pH_med_), and the Donnan Free Space, where electrostatic charges from the wall concentrate cations (including protons) and depletes anions (15). We propose that the proton binding capacity of the cell wall results in pH_apo_ being lower than pH_med_, providing the PM ATPase is sufficently active. In line with this, the apoplast contained within the cell wall was found to be as much as 2 units more acidic than the remaining apoplastic liquid, which was attributed the cell wall’s high cation exchange capacity (16). Moreover, Martinière et al. (17) found that the apoplast is considerably more acidic than the apoplastic liquid immediately adjacent to the PM. Hence, the cell wall may act as a proton reservoir, that charges itself through the action of the PM ATPase and by acquiring protons from the surrounding medium, thus helping to sustain apoplastic acidity. To explain how NRT1.1A/B enhances apoplast acidification, we posit that nitrate symport enhances the activation of H^+^-ATPase, in order to maintain a stable cytosolic pH (Fig. 2*B*).

## Materials and Methods

The detailed methods are described in SI Appendix. Briefly, *M. truncatula* seedlings were grown on agar plates containing nitrate-free FP nutrient solution for one week. The seedlings were then transplanted into non-aerated liquid FP medium supplemented with different concentrations of KNO_3_ with their roots fully submerged and pH was monitored using a pH meter. For HPTS staining, the seedlings were prepared in the same manner and then transferred to FP medium containing 5 mM KNO_3_ for the given time, and then stained with HPTS for 30 minutes, rinsed three times with FP, then root epidermal cells near the root tip were imaged using a confocal microscope. The *M. truncatula nrt1*.*1* and *nrt2*.*1* mutants were previously described (7, 18).

## Acknowledgments

We would like to acknowledge Ping Xu, Yisheng Wang and Chao-Feng Huang for their comments on the manuscript and Wenxian Lan and Wenjuan Cai of the CEMPS core facility for technical support. Seeds of the *nrt2*.*1* mutant were kindly provided by Dr. Fang Xie.

## Author Contributions

QX, JW, YC and WJ performed experimental work and data analysis. JDM and AM supervised and co-directed the research. QX, AJM, FL and JDM conceived the experiments and wrote the article.

## Extended methods

### Seed sterilization

Wild type *M. truncatula* (ecotype R108) and *nrt1*.*1a, nrt1*.*1b, nrt1*.*1c, nrt2*.*1* and the *nrt1*.*1a/b* double mutant seeds were scarified with sandpaper, then surface-sterilized with 70% ethanol for 1 minute, then 10% bleach for 3 minutes, and then rinsed five times with sterile distilled water.

### Plant culture

Sterilized *M. truncatula* (ecotype R108) and *nrt1*.*1a, nrt1*.*1b, nrt1*.*1c, nrt2*.*1* and the *nrt1*.*1a/b* double mutant seeds were placed on 0.8 % agar plates and stratified in darkness at 4°C for 3 days. The plates containing the seedlings were then placed upside down in the dark at 22°C to allow the roots to extend overnight. The seedlings were then transferred to fresh 0.8 % agar plates containing Fahraeus nitrogen free nutrient medium (FP) for one week with the plates vertically positioned.

### pH measurements

For the pH measurements five of the seedlings were transferred to graduated 15 ml plastic tubes filled to the 15 ml mark with FP medium (∼13.2 ml of FP) supplemented with different concentrations of nitrate with the pH adjusted using H_2_SO_4_ or KOH. Plants were grown under controlled conditions with a 16-hour light period at 22°C and an 8-hour dark period at 20°C. The pH of the liquid medium (pH_med_) was measured every few hours using a pH meter (Mettler Toledo FE28) with a reported accuracy of ± 0.01.

### HPTS staining and imaging

HPTS is a membrane impermeant pH sensitive fluorescent dye^1,2,3^. For HPTS staining, plants were grown on FP medium for one week as described above and then transferred to FP medium containing 5 mM KNO_3_ for 0, 1.5, 4, 24, or 48h. Seedlings were then transferred to the same medium supplemented with 1 mM 8-hydroxypyrene-1,3,6-trisulfonic acid (HPTS) (Abcam, USA) for 30 minutes. Roots were then rinsed three times with HPTS-free growth medium, placed on a microscope slide, and covered with a coverslip. Imaging of epidermal cells near the root tip was carried out using an inverted Leica SP8 confocal microscope with a 20X water immersion objective.

HPTS fluorescence was detected using dual excitation at 405 nm and 458 nm, with emission collected at 514 nm. The overall 458/405 ratio was calculated for each image using Fiji software.

### Seed sterilization

Wild type *M. truncatula* (ecotype R108) and *nrt1*.*1a, nrt1*.*1b, nrt1*.*1c, nrt2*.*1* and the *nrt1*.*1a/b* double mutant seeds were scarified with sandpaper, then surface-sterilized with 70% ethanol for 1 minute, then 10% bleach for 3 minutes, and then rinsed five times with sterile distilled water.

### Aeration experiments

To provide gentle aeration of the medium a perforated plastic tube (0.5 mm outer diameter) was placed in each 15 ml tube and connected to an SB-748 silent air pump (Zhongshan SOBO electric appliance co., China) on its lowest setting.

### Measurement of oxygen levels

Plants were grown on FP medium for one week as described above and then transferred to aerated or non-aerated FP or FP supplemented with 5 mM KNO_3_ and left for approximately 30 hours. Oxygen concentrations were then measured using an optical oxygen microsensor (PreSens; Regensburg, Germany).

### PS-1 treatment

PS-1 (TargetMol Chemicals Inc., USA) was stored as a 25 mM stock solution in DMSO. One-week-old seedlings were prepared as described above were transferred to 15 mL tubes containing aerated FP solution supplemented with PS-1 or mock treatment (equivalent amount of 25 µM DMSO) for 6 hours. The pH of the liquid medium was then measured.

### Statistical analyses

All means were compared using Student’s *t*-test or, when more than two means were compared to controls using Dunett’s Test to correct for multiple comparisons.

## References

1. H.-H. Tsai, and W. Schmidt, The enigma of environmental pH sensing in plants. Nat. Plants 7:106–115 (2021).

2. A.J. Miller, and S.J. Smith, Nitrate transport and compartmentation in cereal root cells. J. Exp. Bot. 47: 843–54 (1996).

3. M. Cerezo, et al., Major alterations of the regulation of root NO_3_-uptake are associated with the mutation of Nrt2.1 and Nrt2.2 genes in Arabidopsis. Plant Phys. 127:262–71 (2001).

4. Y.F. Tsay, J.I. Schroeder, K.A. Feldmann, N.M. Crawford, The herbicide sensitivity gene CHL1 of Arabidopsis encodes a nitrate-inducible nitrate transporter. Cell 72:705–13 (1993).

5. A. Maghiaoui, A. Gojon, L. Bach, NRT1.1-centered nitrate signaling in plants. J. Exp. Bot. 71:6226–6237 (2020).

6. X.Z. Fang et al., Alleviation of proton toxicity by nitrate uptake specifically depends on nitrate transporter 1.1 in Arabidopsis. New Phyt. 211:149–58 (2016).

7. Q. Xiao, et al., MtNPF6.5 mediates chloride uptake and nitrate preference in Medicago roots. EMBO J. 40(21):e106847 (2021).

8. M. Haruta et al., Molecular characterization of mutant Arabidopsis plants with reduced plasma membrane proton pump activity. J. Biol. Chem. 285:17918–29 (2010).

9. J.Y. Pang, I. Newman, N. Mendham, M. Zhou, S. Shabala, Microelectrode ion and O_2_ fluxes measurements reveal differential sensitivity of barley root tissues to hypoxia. Plant, Cell & Environ. 29:1107–21 (2006).

10. P.R. McClure, L.V. Kochian, R.M. Spanswick, J.E. Shaff, Evidence for cotransport of nitrate and protons in maize roots: I. Effects of nitrate on the membrane potential. Plant Phys. 93:281–9 (1990).

11. S. Santi, G. Locci, R. Pinton, S. Cesco, Z. Varanini, Plasma Membrane H+-ATPase in maize roots induced for NO_3_^-^uptake. Plant Phys. 109:1277–1283 (1995).

12. S. Santi, G. Locci, R. Monte, R. Pinton, Z. Varanini, Induction of nitrate uptake in maize roots: expression of a putative high-affinity nitrate transporter and plasma membrane H+-ATPase isoforms. J. Exp. Bot. 54:1851–64 (2003).

13. L. Espen, F.F. Nocito, M. Cocucci, Effect of NO_3_^-^transport and reduction on intracellular pH: an in vivo NMR study in maize roots. J. Exp. Bot. 55:2053–61 (2004).

14. H. Sze, J.M. Ward, S. Lai, Vacuolar ^H+^-translocating ATPases from plants: structure, function, and isoforms. J Bioenerg. and Biomemb. 24:371–81 (1992).

15. D. S. Bush, and J. G. McColl, Mass-action expressions of ion exchange applied to Ca^2+^, H^+^, K^+^, and Mg^2+^ sorption on isolated cell walls of leaves from Brassica oleracea. Plant Physiol. 85:247–60 (1987).

16. H. Sentenac, C. Grignon, Effect of H^+^ excretion on the surface pH of corn root cells evaluated by using weak acid influx as a pH probe. Plant Phys. 84:1367–1372 (1987).

17. A. Martinière et al., Uncovering pH at both sides of the root plasma membrane interface using noninvasive imaging. Proc. Nat. Acad. Sci. U.S.A. 115:6488–6493 (2018).

18. Z. Luo, et al., The small peptide CEP1 and the NIN-like protein NLP1 regulate NRT2.1 to mediate root nodule formation across nitrate concentrations. Plant Cell 35:776–794 (2023).

## References

1. Lin W, Zhou X, Tang W, Takahashi K, Pan X, Dai J, Ren H, Zhu X, Pan S, Zheng H, et al. 2021. TMK-based cell-surface auxin signalling activates cell-wall acidification. Nature 599: 278–282.

2. Wang J, Jin D, Deng Z, Zheng L, Guo P, Ji Y, Song Z, Zeng HY, Kinoshita T, Liao Z, et al. 2025. The apoplastic pH is a key determinant in the hypocotyl growth response to auxin dosage and light. Nature Plants 11:279–294.

3. Barbez E, Dünser K, Gaidora A, Lendl T, Busch W. 2017. Auxin steers root cell expansion via apoplastic pH regulation in Arabidopsis thaliana. Proc. Natl. Acad. Sci. U.S.A. 114(24):E4884–E4893.

